# TALE: Transformer-based protein function Annotation with joint sequence–Label Embedding

**DOI:** 10.1101/2020.09.27.315937

**Authors:** Yue Cao, Yang Shen

## Abstract

**Motivation:** Facing the increasing gap between high-throughput sequence data and limited functional insights, computational protein function annotation provides a high-throughput alternative to experimental approaches. However, current methods can have limited applicability while relying on data besides sequences, or lack generalizability to novel sequences, species and functions.

**Results:** To overcome aforementioned barriers in applicability and generalizability, we propose a novel deep learning model, named Transformer-based protein function Annotation through joint sequence–Label Embedding (TALE). For generalizbility to novel sequences we use self attention-based transformers to capture global patterns in sequences. For generalizability to unseen or rarely seen functions, we also embed protein function labels (hierarchical GO terms on directed graphs) together with inputs/features (sequences) in a joint latent space. Combining TALE and a sequence similarity-based method, TALE+ outperformed competing methods when only sequence input is available. It even outperformed a state-of-the-art method using network information besides sequence, in two of the three gene ontologies. Furthermore, TALE and TALE+ showed superior generalizability to proteins of low homology and never/rarely annotated novel species or functions compared to training data, revealing deep insights into the protein sequence–function relationship. Ablation studies elucidated contributions of algorithmic components toward the accuracy and the generalizability.

**Availability:** The data, source codes and models are available at https://github.com/Shen-Lab/TALE

**Supplementary information:** Supplementary data are available at *Bioinformatics* online.

## 1 Introduction

The explosive growth of protein sequence data in the past few decades, largely thanks to next-generation sequencing technologies, has provided enormous information and opportunities for biological and pharmaceutical research. In particular, the complex and intricate relationship between sequences, structures, and functions of proteins is fascinating. As experimental function annotation of proteins is often outpaced by sequence determination, computational alternatives have become both fundamental in exploring the sequence-function relationship and practical in predicting functions for growing un-annotated sequences (including *de novo* designs). According to the 2020_01 release of UniProt (UniProtConsortium, 2019), there were around 5.6×10^5^ nonredundant sequences manually annotated in Swiss-Prot but over two orders of magnitude more (around 1.8×10^8^) sequences awaiting full manual annotation in TrEMBL.

Protein functions are usually described based on Gene Ontology (GO), the world’s largest source of systematic representation of gene functions (Ashburner *et al.*, 2000). There are more than 40,000 GO terms across three domains: Molecular Function Ontology (MFO), Biological Process Ontology (BPO) and Cellular Component Ontology (CCO). Within each ontology, GO terms are structured hierarchically as a directed acyclic graph (DAG) with a root node. Each protein can be annotated with more than one GO term on three ontologies (thus a multi-label classification problem). If a protein is annotated with one GO term, then can also be annotated by all corresponding ancestral GO terms. Such a hierarchical constraint is present in many other ontologies as well, such as text ontology (Baker and Korhonen (2017)) and image ontology (Deng *et al.* (2009)).

From the perspective of input type, computational methods for protein function annotation can be classified as sequence- (Fa *et al.*, 2018; Kulmanov and Hoehndorf, 2020; Zhou *et al.*, 2019a), structure- (Yang *et al.*, 2015), network- (You *et al.*, 2019; Kulmanov *et al.*, 2018), and literature-based (Kahanda and Ben-Hur, 2017; You *et al.*, 2018a), whereas all but sequence-only methods have limited scope of usage due to data availability. Specifically, although structural information is important for understanding protein functions (e.g. (Stewart *et al.*, 1998; Wrapp *et al.*, 2020)), it is often not readily available: 0.01% of the TrEMBL sequences (UniProt 2020_01) have corresponding structural entries in Protein Data Bank (PDB), and this ratio increases to 1% if considering structural models in SWISS-MODEL Repository (SWR). Similarly, only around 7% of the TrEMBL sequences have interaction entries in STRINGdb (Szklarczyk *et al.*, 2016), not to mention that the network information can be noisy and incomplete. We note that structure, network, and literature data can be especially missing for novel sequences where computational function annotation is needed the most. We therefore focus on sequence-based methods in this study.

Predicting protein function from sequence alone is a challenging problem where each sequence can belong to multiple labels and labels are organized hierarchically. Critical Assessment of protein Function Annotation (CAFA) has provided an enabling platform for method development (Radivojac *et al.*, 2013; Jiang *et al.*, 2016; Friedberg and Radivojac, 2017; Zhou *et al.*, 2019b) and witnessed still-limited power or scope of current methods. Sequence similarity-based methods (Jones *et al.*, 2005; Buchfink *et al.*, 2015) leverage sequence homology, although their success is often limited to homologues and alignments to detect homology can be still costly. Recently, deep learning has emerged as a promising approach (Kulmanov *et al.*, 2018; Kulmanov and Hoehndorf, 2020) to improve the accuracy, where sequences are often inputs/features and GO terms are labels. However, as deep learning is a data-hungry technique, these methods often have to get rid of a large number of GO terms (labels) with few annotations, leading to narrow applicability. For instance, DeepGOPlus (Kulmanov and Hoehndorf, 2020) only considered over 5,000 GO terms with at least 50 annotated sequences each, which only accounts for less than 12% of all GO terms.

We set out to overcome aforementioned barriers and boost the generalizability to low sequence homology as well as unseen or rarely seen functions (also known as tail labels). To that end, we propose a novel approach named Transformer-based protein function Annotation through joint sequence–Label Embedding (TALE). Our contributions are as follows. First, TALE replaces previously-used convolutional neural networks (CNN) with self-attention-based transformers (Vaswani *et al.*, 2017) which has made a major breakthrough in natural language processing and recently in protein sequence embedding (Rives *et al.*, 2019; Duong *et al.*, 2020; Elnaggar *et al.*, 2020). Compared to CNN, transformers can deal with global dependencies within the sequence in just one layer, which helps detect global sequence patterns for function prediction much easier than CNN-based methods do. Second, TALE embeds sequence inputs/features and hierarchical function labels (GO terms) into a latent space. By considering similarities among function labels and sequence features, TALE can easily deal with tail labels. Unlike previous methods that only consider GO terms as flat labels and enforce hierarchy *ad hoc* after training, TALE considers the hierarchy among labels through regularization during training. Last, we propose TALE+, by using an ensemble of top three TALE models and a sequence similarity-based method, DIAMOND (Buchfink *et al.*, 2015), in convex combination (similar to DeepGoPlus) to reach the best of both worlds.

Over all Swiss-Prot sequences experimentally-annotated and released since 2017, we have empirically compared TALE and TALE+ to six baseline methods including NetGO (You *et al.*, 2019), the upgraded version of GoLabeler (You *et al.*, 2018b), the top performer in the latest CAFA. TALE+ outperforms all other methods in all three ontologies when only sequence information is used. Importantly, TALE and TALE+ significantly improve against state-of-the-art methods in challenging cases where test proteins are of low similarity (in sequence, species, and label) to training examples. The results prove that our model can generalize well to novel sequences, novel species and novel functions.

The rest of the paper is organized as follows. We will first discuss in Methods the data set used in the study. We will then introduce TALE and TALE+ models in details besides baselines and end the section with evaluation metrics. We will start the Results section with overall performance comparison. We will then delve into the analysis on model generalizability in sequence, species and function. Lastly, we will report an ablation study to delineate the major algorithm contributors to TALE’s improved generalizability.

## 2 Methods

### 2.1 Datasets

We use functionally annotated protein sequences from UniProtKB/Swiss-Prot and hierarchical relationships between function labels (GO terms) from Gene Ontology.

#### 2.1.1 Functionally-annotated Protein Sequences

We downloaded 561,176 protein sequences from UniProtKB/Swiss-Prot 2019_09. We filtered these sequences based on the following criteria: (1) sequences whose lengths are above 1,000 or who contain non-standard amino acids were removed, which constitute 3.3% of all sequences; (2) consistent with Critical Assessment of protein Function Annotation (CAFA) (Zhou *et al.*, 2019b), only sequences with high-quality function annotations were retained, i.e., those with at least one annotation within the following 8 experimental evidence codes: EXP, IDA, IPI, IMP, IGI, IEP, TAS, and IC.

#### 2.1.2 Time-split Datasets

We split the resulting dataset according to the time when sequences were released in Swiss-Prot. Specifically, we extract the sequences by the end of 2012 to be the training set, those between 2013 and 2016 to be the validation set, and those since 2017 to be the test set. The statistics of sequences and GO terms over these datasets and three ontologies are shown in Table 1. We will train all models on the training set and tune their hyperparameters with the validation set. And finally we will retrain models under their optimal hyperparameters, using both the training and the validation sets, and assess them on the test set.

**Table 1.** Statistics of sequences and GO terms in various ontologies and datasets.

| Ontology | MFO | BPO | CCO |
| --- | --- | --- | --- |
| #Seq in Training Set | 30541 | 43107 | 38601 |
| #Seq in Validation Set | 2454 | 3492 | 2139 |
| #Seq in Train. + Val. | 32995 | 46599 | 40740 |
| #Seq in Test Set | 1559 | 2610 | 1907 |
| #Seq in Test Set without network information | 465 | 726 | 511 |
| #GO terms | 6421 | 20673 | 2677 |

#### 2.1.3 Hierarchical Relationships between GO Terms

We downloaded the identities and the hierarchical relationships between function labels (GO terms) from Gene Ontology (Ashburner *et al.*, 2000) (data-version: releases/2020-01-01). We consider ‘is a’ and ‘part of’ relationships in all three ontologies: Molecular Function Ontology (MFO), Biological Process Ontology (BPO) and Cellular Component Ontology (CCO); and do not consider cross-ontology relationships for now. In this way we had three separate ontologies, each of whose topology is a directed acyclic graph (DAG). For each annotation in each ontology, we additionally propagated annotations all the way from the corresponding GO term to the root node. Lastly, 14,929 out of 44,700 GO terms without a single annotated sequence were removed and not considered in this study.

### 2.2 TALE and TALE+

We will describe the details of our methods in this subsection. The overall architecture of TALE is shown in Fig. 1. The model has two inputs: a protein sequence and the label matrix (for capturing hierarchical relationships between all GO terms). It is worth mentioning that the label matrix is a constant matrix for a given ontology, thus being fixed during both stages of training and inference. The model itself consists of feature (sequence) embedding, label (function) embedding, joint similarity modules, as well as fully-connected and output (softmax) layers. We will introduce these components in the following subsections.

**Fig. 1:**
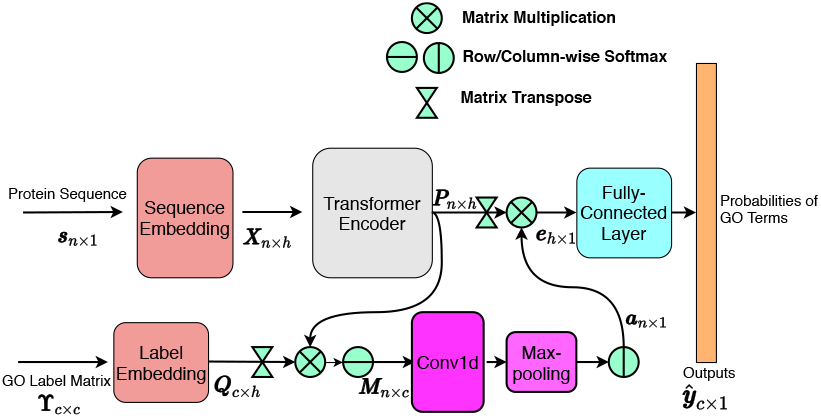
The architecture of TALE. Note that the GO label matrix is fixed for each ontology.

*Notations*. We use upper-case boldfaced letters to denote a matrix (e.g. ***X***), lower-case boldfaced letters to denote a vector (e.g. ***x***), and lower-case letters to denote a scalar (e.g. *x*). We use subscripts of a matrix to denote a specific row, column, or element (e.g. ***X***_*i*_ for the *i*th row of ***X***; ***X***_,*i*_ for the *i*th column; and ***X***_*i,j*_ for the entry in the *i*th row and the *j*-th column of ***X***). We also use subscripts to denote scalar components of a vector (e.g. *x*_*i*_ for the *i*th entry of the vector ***x***). We use superscript *T* on a matrix to represent its transpose.

#### 2.2.1 Sequence Embedding

Let 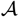 denote the set of 20 standard amino acids plus the padding symbol. For a given input protein sequence 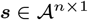 of length *n*, we embed each character (residue) into an *h*-dimensional continuous latent space through a trainable lookup matrix ***W***^seq^ and positional embedding, as described in (Vaswani *et al.*, 2017). The embedded matrix ***X*** ∈ ℝ^*n*×*h*^ is fed through a transformer encoder that consist of multiple multi-head attention layers. In this study, we used 6 layers and 2 heads for each layer. The advantage of such “self-attention” layers compared to convolution layers is that self-attention can easily and quickly capture long-term dependencies within a whole sequence, whereas convolution can only capture dependencies of residues within a neighborhood determined by convolutional kernals. We denote the output matrix of the transformer encoder with ***P*** ∈ ℝ^*n*×*h*^, where *h* is the hidden dimension.

#### 2.2.2 Label Embedding

For each given ontology represented as a directed acyclic graph (DAG), we first perform topological sorting of its nodes (GO terms or labels) and assign an index to each node based on its order in the sorted array. We then embed node *i* into a *c*-dimensional binary vector 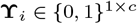 where *c* is the number of labels or GO terms. In this way we embed all nodes within the ontology with a label matrix 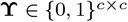 where its *i*th row 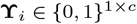 is the embedding vector for node *i* and its element 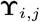 is 1 if node *j* is an ancestor of node *i* and 0 otherwise: 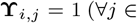 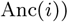, where Anc(*i*) denotes the set containing all ancestors of node *i* (plus *i* itself). Similar to the sequence embedding, we use a trainable lookup matrix ***W***^label^ ∈ ℝ^*c*×*h*^ to encode 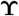 as ***Q*** ∈ ℝ^*c*×*h*^, where

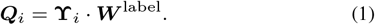

Unlike sequence input, the label matrix 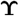 is fixed for each ontology and thus not needed to be further encoded using a transformer encoder.

#### 2.2.3 Joint Sequence–Label Similarity

We inspect the contributions of individual amino acids to individual function labels, by calculating the matrix product between ***P*** and ***Q*** to measure the joint similarity between the sequence and the label:

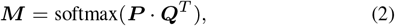

where the softmax is row-wised. ***M***_*i,j*_ suggests the “closeness” or similarity score between amino acid *i* and label *j*. For each amino acid *i*, we further consider the contributions from other amino acids, by applying a 1D convolutional layer to ***M*** (along the row direction with the columns as channels), followed by a max-pooling layer. The output of the max-pooling layer is first normalized, and then used for weighting the sequence encoding matrix ***P***:

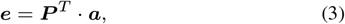

where ***a*** ∈ ℝ^*n*×1^ is the output of the max-pooling after column-wise softmax.

#### 2.2.4 Fully-connected and Output Layers

The output of the joint similarity module, ***e*** ∈ ℝ^*h*×1^, would go through two fully-connected (FC) layers, with the sigmoid activation function at the second FC layer. The output of the model 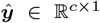 is the predicted probabilities for individual GO terms in the ontology, where the *i*th component is the predicted probability of label *i* for a given input sequence.

#### 2.2.5 Loss and Hierarchical Regularization

To train model parameters, we first consider the binary cross-entropy loss:

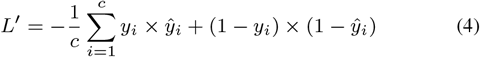

However, if we only use *L*′, trained models may make predictions violating the hierarchical constraint of function annotation. For instance, the predicted score (probability) of a child GO term may be larger than the scores of ancestors. To mitigate such hierarchical violation, we then introduce an additional, hierarchical regularization term:

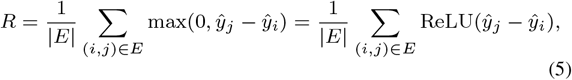

where *E* is the set of all edges in the ontology graph, and (*i, j*) is one edge in *E* pointing from node *i* to *j*. Therefore, our overall loss function is a weighted sum of both terms: *L* = *L*′ + *λR*, where *λ*, the regularization constant to control the balance between the two terms, is treated as a hyper-parameter and tuned along with other hyper-parameters using the validation set.

#### 2.2.6 Ensemble Model of TALE+

So far we have introduced all components of TALE. In order to reduce the variance of predicted scores and their generalization errors, we consider to first use the simple average of the outputs of top 3 models based on the validation set, as the final TALE predictions.

Similar to DeepGOPlus (Kulmanov and Hoehndorf, 2020), we further use a convex combination of TALE (the simple average) and DIAMONDScore as final outputs of TALE+:

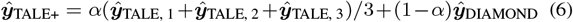

where 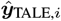 is the *i*th best TALE model based on the validation set. The DIAMONDScore method will be introduced in **Sec**. 2.3.2. After tuning on the validation set, the best *α*s for three ontologies were set to be 0.4 for MFO, 0.5 for BPO, and 0.7 for CCO. TALE can be regarded as a special case of TALE+ when *α* = 1.

### 2.3 Baseline and State-of-the-Art Methods

We compare TALE and TALE+ to six competing methods, including baselines, latest published methods and top performers in CAFA.

#### 2.3.1 Naive Approach

One simple approach is to use the background frequency of individual GO terms in the training set to annotate every query sequence: 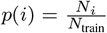, where *i* is the *i*th GO term, *N*_*i*_ is the number of sequences in the training set annotated with the *i*th GO term, and *N*_train_ is the total number of sequences in the training set. This simple approach is called “naive” approach in CAFA and is used as the baseline approach for comparing methods.

#### 2.3.2 DIAMONDScore

DIAMONDScore is a sequence similarity-based function annotation method. For a query sequence ***q***, we use DIAMOND (Buchfink *et al.*, 2015) to find its similar sequences (homologs) in the training set and obtain a bitscore for each pair of query sequence and a homolog in the training set: bitscore(***q***, ***s***), 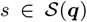, where 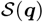 denotes the set of training sequences that are similar to the query sequence under an E-value cutoff of 0.001. Then for the *i*th GO term GO_*i*_, we calculate the predicted probability associated with ***q*** as:

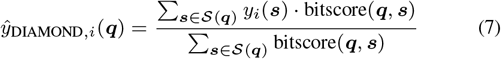

where *y*_*i*_(***s***) is the label of *i*th GO term for the training sequence ***s***.

#### 2.3.3 DeepGO, DeepGOCNN and DeepGOPlus

DeepGO (Kulmanov *et al.*, 2018) is a deep learning method that uses 1D convolutional networks on protein sequences for structural outputs (i.e. hierarchical classification of protein functions). In addition, DeepGO use protein-protein interaction (PPI) features.

DeepGOCNN and DeepGOPlus are recently published improvements (Kulmanov and Hoehndorf, 2020). Compared to DeepGO, DeepGOCNN ignores the PPI features, trainable embedding and structural outputs. In this way, DeepGOCNN is more efficient and applicable for a much larger portion of proteins and GO terms. Then DeepGOPlus combines the outputs of DeepGOCNN and those of DIAMONDScore through convex combination.

#### 2.3.4 NetGO

NetGO (You *et al.*, 2019) is a hybrid method, which merges the network information into its previous sequence-based method GOLabeler (You *et al.*, 2018b). GOLabeler is the winner of CAFA3 and uses a “learning to rank” framework to rank GO terms.

### 2.4 Implementation

Our models were implemented in Tensorflow 1.13 (Abadi *et al.*, 2016) and trained for 100 epochs using Adam (Kingma and Ba, 2014) on a single Nvidia Tesla K80 GPU. The hyperperameters were tuned on the validation set mainly based on Fmax, a major assessment metric used in CAFA (see Sec.2.5 for definition). AuPRUC was used as a tie-breaker when Fmax values are within 0.003. The list of hyperparameters and their optimal values are provided in the Supplemental Table S1. The curves for training loss and validation accuracy of the best single TALE model are shown in Figure S1 for each ontology.

For DeepGO, DeepGOCNN and DeepGOPlus, we used their published codes on Github and trained the models on our datasets. For DeepGOCNN and DeepGOPlus, we tuned the hyperparameters on our validation set. Specifically, we set the ‘max_kernel’ to be 64, and ‘nb_filters’ to be 256. The learning rate of Adam optimization algorithm was set to be 1e-3. The constant *α* after tuning was 0.5 in MFO, 0.6 in BPO, and 0.6 in CCO. We have also shown the number of trainable parameters of DeepGOPlus (same as DeepGOCNN) against TALE+ in Table S2. It can be seen that the amount of trainable parameters in TALE (considering top 3 models for each of the three ontologies, i.e., 9 models in total) is below 40% of DeepGOPlus.

For NetGO, we only have access to its webserver.

### 2.5 Evaluation

For a test set *D*_test_, we choose to use two evaluation metrics: Fmax and AuPRC. Fmax is the official, protein-centric evaluation metric used in CAFA. It is the maximum score of the geometric average of averaged precision and recall over proteins for all thresholds:

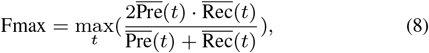

where 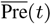 and 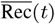 are the averaged precision and recall at threshold *t*. Specifically,

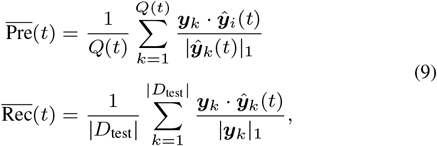

where *Q*(*t*) is the number of samples which contain at least one non-zero label in *D*_test_; ***y***_*i*_ is the true label vector of *k*th sample in *D*_test_, and 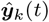 is the predicted label vector of the *k*th sample in *D*_test_ at threshold *t*. For the calculation, we iterated *t* incrementally from 0 to 1 at a stepsize of 0.01.

AuPRC is a standard metric in machine learning for evaluating the binary classification performance, especially suitable for highly imbalance data, which is often the case in protein function annotation. In multi-label classification, we concatenate all the label vectors and use canonical AuPRC (single-label) to evaluate the performance.

## 3 Results

We perform comprehensive evaluation of our models from several perspectives. We will start with comparing them to aforementioned competing methods on various ontologies and test sets. We will proceed to assess the capability of all models to generalize to novel sequences, novel species, and novel functions relative to the training set. And we will conclude with an ablation study for TALE and TALE+ to delineate the contributions of their various algorithm components to performances and generalizability.

### 3.1 Performance on the test set

We compare TALE and TALE+ with competing methods over the test set and show the results in Table 2. Overall, TALE+ achieved the best performance in biological process (BPO) and cellular component (CCO); and had the second best in molecular function (MFO). The best performer in MFO, NetGO was also the second best performer in BPO and the third in CCO. Specifically, compared to NetGO, TALE+ improved Fmax from 0.425 to 0.438 (by 3%) in BPO and from 0.672 to 0.728 (by 8%) in CCO; and it had worse Fmax (0.679 vs. 0.691) in MFO. TALE alone without adding similarity-based DIAMONDScore outperformed all other methods except our own TALE+ in CCO: compared to NetGO, TALE alone improved Fmax from 0.672 to 0.699 (by 4%). It is noteworthy that NetGO uses additional network information that is not used in TALE or TALE+. Such network information is often not available to proteins and, in such cases, TALE+ outperformed NetGO in all three ontologies including MFO, as will be shown in Table 3,

**Table 2.** The performance of TALE and TALE+ against competing methods on the test set. Note that both DeepGO and NetGO use network information besides sequence, whereas other methods including TALE and TALE+ use sequence alone.

| Ontology | Fmax |  |  | AuPRC |  |  |
| --- | --- | --- | --- | --- | --- | --- |
|  | MFO | BPO | CCO | MFO | BPO | CCO |
| Naive | 0.399 | 0.278 | 0.634 | 0.320 | 0.194 | 0.589 |
| DIAMONDScore | 0.600 | 0.377 | 0.564 | 0.533 | 0.249 | 0.476 |
| DeepGO | 0.432 | 0.258 | 0.587 | 0.355 | 0.201 | 0.516 |
| DeepGOCNN | 0.458 | 0.310 | 0.599 | 0.345 | 0.176 | 0.506 |
| DeepGOPlus | 0.635 | 0.398 | 0.661 | 0.562 | 0.258 | 0.590 |
| NetGO | **0.691** | 0.425 | 0.672 | **0.615** | **0.316** | 0.642 |
| TALE | 0.585 | 0.363 | 0.699 | 0.564 | 0.280 | 0.685 |
| TALE+ | 0.679 | **0.438** | **0.728** | 0.613 | **0.316** | **0.711** |

**Table 3.** The performance of TALE+ against competing methods on the portion of test set that does not have network information available.

| Ontology | Fmax |  |  | AuPRC |  |  |
| --- | --- | --- | --- | --- | --- | --- |
|  | MFO | BPO | CCO | MFO | BPO | CCO |
| Naive | 0.412 | 0.267 | 0.558 | 0.320 | 0.183 | 0.544 |
| DIAMONDScore | 0.603 | 0.344 | 0.549 | 0.545 | 0.233 | 0.438 |
| DeepGO | 0.413 | 0.266 | 0.576 | 0.343 | 0.223 | 0.507 |
| DeepGOCNN | 0.459 | 0.307 | 0.611 | 0.342 | 0.181 | 0.470 |
| DeepGOPlus | 0.631 | 0.372 | 0.660 | 0.569 | 0.255 | 0.524 |
| NetGO | 0.645 | 0.375 | 0.642 | 0.605 | 0.274 | 0.531 |
| TALE | 0.538 | 0.343 | 0.746 | 0.481 | 0.243 | 0.682 |
| TALE+ | **0.671** | **0.416** | **0.756** | **0.613** | **0.291** | **0.687** |

The performance difference among the three ontologies can be attributed to the structure and complexity of the ontology as well as the available annotations (Jiang *et al.*, 2016). Similarity-based DIAMONDScore performed much better than the naive approach (and well among all methods) in MFO but worse in CCO. This observation echos the hypothesis that sequence similarity may carry more information on basic biochemical annotations than cellular components. Interestingly, CCO is also the ontology where TALE and TALE+ did the best – even TALE alone was better than all methods other than TALE+ and adding similarity-based DIAMONDScore to TALE resulted in less help than it did in MFO and BPO.

### 3.2 Performance on the test set without network information

Protein-protein interaction (PPI) network information can be very useful in boosting the accuracy of computational protein function annotation. However, its availability can be limited and, when available, its quality can also be limited (noisy and incomplete). Table 1 shows that 70% of our test set has corresponding network information in STRINGdb, which is already biased considering that all test sequences have already been functionally annotated with experiments until now. In reality, only 7% of TrEMBLE sequences have corresponding STRINGdb entries available, regardless of the quality of the network information. The ratio can be even worse for *de novo* designed protein sequences. Therefore, a reliable sequence-only function annotation method is necessary especially when network information is not available.

We thus did performance analysis over the portion of the test set without network information, i.e, the test sequences whose network information cannot be found in STRINGdb (see statistics in Table 1). In this case alone we used the version of DeepGO codes without network features. As shown in Table 3, TALE+ significantly outperformed all competing methods in all three ontologies. Compared to NetGO, TALE+ improved Fmax from 0.645 to 0.671 (by 4%) in MFO, from 0.375 to 0.416 (by over 10%) in BPO, and from 0.642 to 0.756 (by nearly 18%) in CCO.

### 3.3 Generalizability from the training set to the test set

Despite the improved performances of our models, questions remain on their practical utility. Are they useful in cases where function annotation is needed the most? Sequence-based protein property prediction (e.g., fold, structure, and function) is needed the most in the “midnight” zone where similarity to known annotated sequences is too low to sustain the assumption that similar inputs (sequences) imply similar outputs (aforementioned properties). Similarly, it is also needed the most in the midnight zone in another sense, where examples for some specific function labels are never or rarely seen in known annotated sequences (thus these functions are referred to as tail labels).

Besides the practical questions on model applicability in the midnight zone, fundamental biological questions also remain. Have these machine learning models learned anything fundamental in structure–function relationships? Or are they merely mimicking patterns in training data using a complicated function (such as a neural network) that could overfit?

To answer the questions above, we examine our models and competing ones in their generalizability from the training set to the test set. Models considered include DIAMOND, DeepGOCNN, TALE and TALE+. (We couldn’t include the NetGO webserver as its training set is not accessible.) Specifically, we examine the generalizability of these models from three perspectives: sequence, species, and label (function) as follows.

#### 3.3.1 Sequence generalizability

To analyze the generalizability to novel sequences, we split the test set into bins of various sequence-identity levels compared to the training set and examine various models’ accuracy (Fmax) over these bins. Specifically, sequence identity between a test sequence and the training set is measured by the maximum sequence identity (MSI). The distributions of test sequences in MSI are shown in Fig. S2. We partition the test sequences into 10 bins equal-spaced in MSI and show the sequence counts (gray dots) and the Fmax scores (colored bars) in these bins (Fig. 2).

**Fig. 2:**
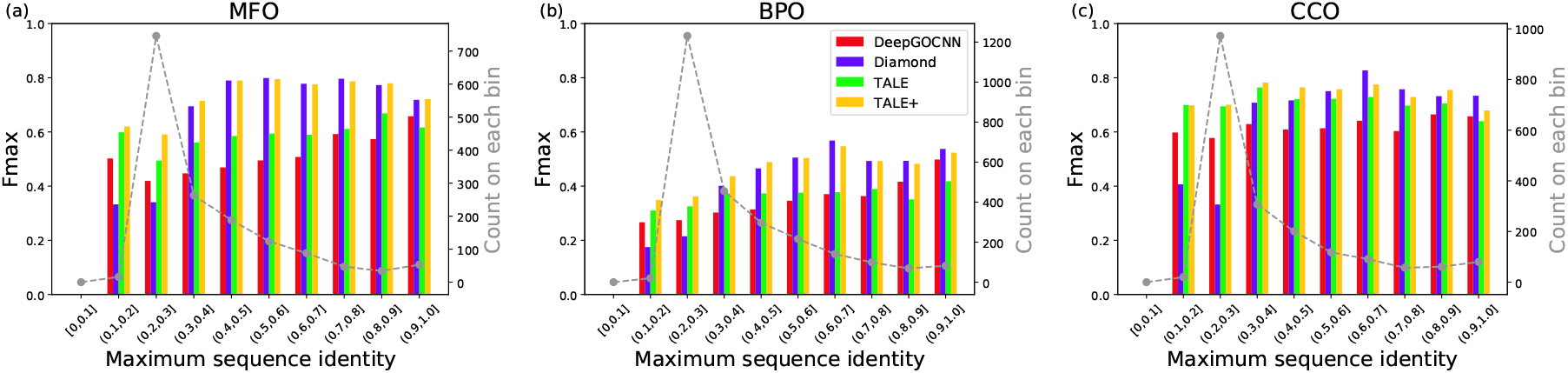
The Fmax performances of four models in three ontologies, over 10 bins of increasing sequence-identity ranges. Low sequence identity indicate low homology between a test sequence and the training set. Sequence statistics over the bins are also provided.

As shown in Fig. 2, sequence similarity-based method DIAMOND performed poorly compared to deep learning-based models, when sequence identity is below 30% (a “midnight” zone). Without a surprise, it was the best or close-to-best performer in MFO, BPO, and CCO, when sequence identity is above 40%, 50%, and 60%, respectively. Between DeepGOCNN and TALE that do not use sequence-similarity scores from other sources, TALE outperformed DeepGOCNN in all bins except above 90% (MFO), 80% (BPO), and 90% (CCO); and it significantly outperformed DeepGOCNN in the “midnight” bins. TALE+ had the best performance in nearly every combination of ontology and bin. As a convex combination of DIAMONDScore and TALE, TALE+ maintained the impressive performances of TALE in the low sequence-identity cases and significantly improved from TALE in the high sequence-identity cases using the similarity-based information.

#### 3.3.2 Species generalizability

To analyze the generalizability to new species, we count for every test sequence the Number of training Samples of the Same Species (NSSS) and identify those in new species never seen in the training set (NSSS=0). The statistics are in Table S2 and the distributions of the test sequences in NSSS are in Fig. S3. We further remove test sequences whose maximum sequence identities to the training set are above 40%. The remaining test sequences are thus in new species and low homology compared to the training set (see statistics in Table S3).

Fig. 3 shows the counts (gray dots) and the Fmax scores of various models (colored bars) over prokaryotes (bacteria and archaea) and eukaryotes. Again, TALE significantly outperformed the similarity-based DIAMOND and deep learning-based DeepGOCNN for nearly all six combinations of clades and ontologies (except being edged by DIAMOND for eukaryotic MFO). Similarity-based DIAMOND performed poorly and provided negligible boost to TALE+ for BPO and CCO in these cases. Interestingly, all methods performed better for new eukaryotic species than for new prokaryotic species in almost every ontology (except for deep learning-based methods in BPO).

**Fig. 3:**
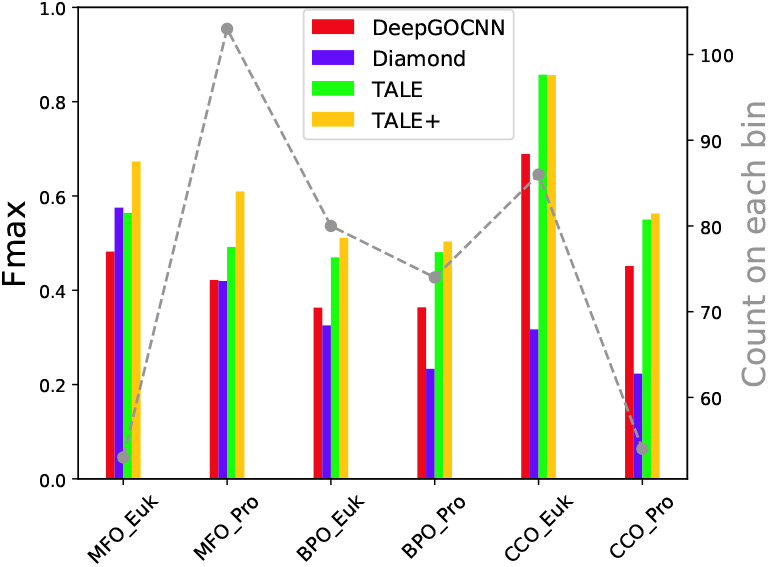
The Fmax performances of four models in three ontologies, over eukaryotes and prokaryotes with NSSS=0 (new species) and MSI ≤ 40% (low homology).

#### 3.3.3 Function generalizability

To analyze the generalizability to new or rarely annotated functions (GO labels), we first calculate the frequency of the *i*th label in the training set:

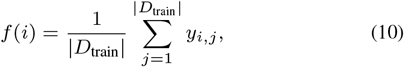

where |*D*_train_| is the number of samples in the training set and the binary *y*_*i,j*_ is for the *i*th label of the *j*th sample in the training set. Then for the *k*th test sample, we calculate its overall label frequency (LF) in the training set as:

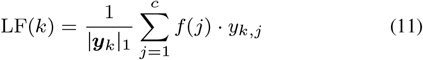

where the binary *y*_*k,j*_ is for the *j*th label of the *i*th sample in the test set, and ***y***_*k*_ is the stacked vector over all labels.

The distributions of test sequences in LF(·) are shown in Fig. S4. We split the test set based into 10 equal-spaced LF(·) bins and provide the histograms of the sequences over these bins in Fig. 4 (gray dots).

**Fig. 4:**
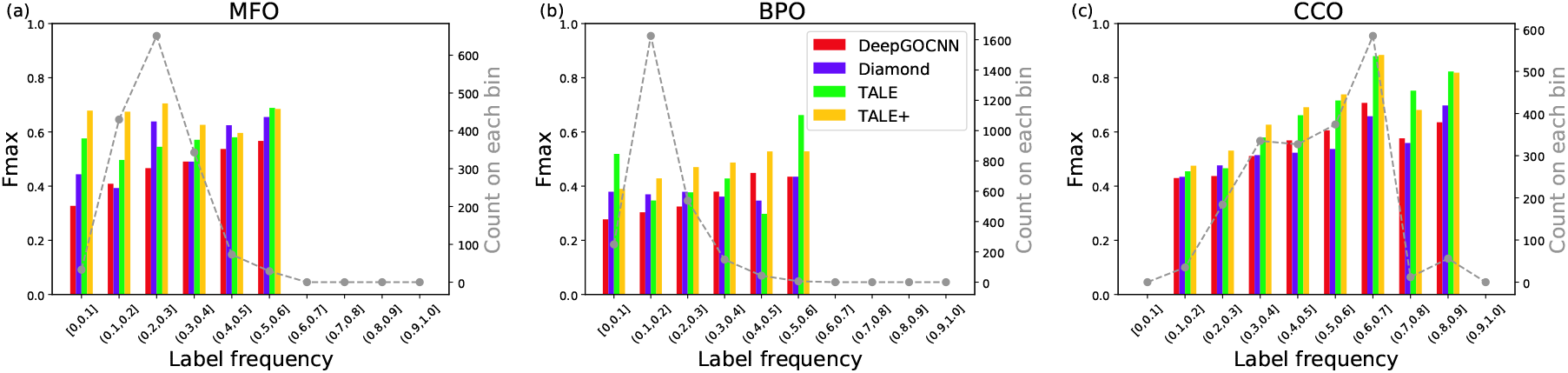
The Fmax performances of four models in three ontologies, over 10 bins of increasing function/label frequencies (for each test sequence, average label frequencies in the training set, measured by LF(·)). Low LF bins indicate functions rarely annotated in the training set.

As shown in Fig. 4, the performances of similarity-based DIAMONDScore, especially in MFO and CCO, did deteriorate noticeably as the (average) label frequency of a test sequence decreases. Interestingly, deep learning alone does not necessarily lead to better performances in such scenarios, as DeepGOCNN actually had worse performances in all ontologies compared to DIAMOND when the test sequences’ function labels are the least frequent in the training set (less than 10%). In this context, the much-improved performances of TALE and TALE+ in the low label-frequency (LF) bins attest to the advantage of our models. Interestingly, adding similarity-based DIAMONDScore to TALE, did not always lead to a further improved TALE+, as seen in BPO (one low and one high LF bins) and CCO (one high LF bin).

In total, similarity-based DIAMOND did not generalize well to novel sequences in low homology (and new species) compared to the training ones, whereas deep learning-based DeepGOCNN performed poorly for never or rarely annotated functions (tail labels). TALE outperformed both methods in all generalizability tests, echoing our earlier rationale that joint embedding of sequences and hierarchical function labels to address their similarities would significantly improve the performance for novel sequences and tail labels. Combining TALE and similarity-based DIAMOND into TALE+ could further enhance the performances in some cases. Generalizability analysis based on equal-populated bins also led to similar conclusions (see details in SI Sec. 6).

### 3.4 Ablation Study

To rigorously delineate the contributions of algorithmic innovations that we have made in TALE and TALE+ to their improved performances and superior generalizability, we perform the following ablation study. Starting with DeepGOCNN, we incrementally add algorithm components and introduce variants to eventually become TALE and TALE+:

- **B1**: replacing the convolutional layers in DeepGOCNN with the transformer encoder plus the input embeddings. Unlike Eq. (3) where ***a*** is the output from joint label embedding, the output of the encoder here ***P*** ∈ ℝ^*n*×*h*^ would be simply row-averaged to obtain ***e***;
- **B2**: replacing the DeepGOCNN post-training correction of hierarchical violations in **B1** with the additional loss term of hierarchical regularization (Eq. 5);
- **B3**: adding label embedding to **B2** for joint sequence–label embedding.

TALE is using the average prediction of the ensemble of top-3 **B3** models (based on the validation set) and TALE+ is a convex combination of TALE and DIAMONDScore.

The overall performances of the above models over the test set are summarized in Fig. 5(a). Exact Fmax and AuPRC values and detailed analyses are in SI Sec. 7. We note that, from DeepGOCNN to TALE+, both Fmax and AuPRC are gradually increasing over model variants. In MFO and BPO, the convex combination with DIAMONDScore (and ensemble average) had the largest contributions to the overall Fmax increases, whereas transformers did so to the AuPRC improvements. In CCO, the transformers, hierarchical regularization, and label embedding (jointly with sequence embedding) played the central role in improving both measures, while similarity-based DIAMONDScore also helped Fmax but not AuPRC.

**Fig. 5:**
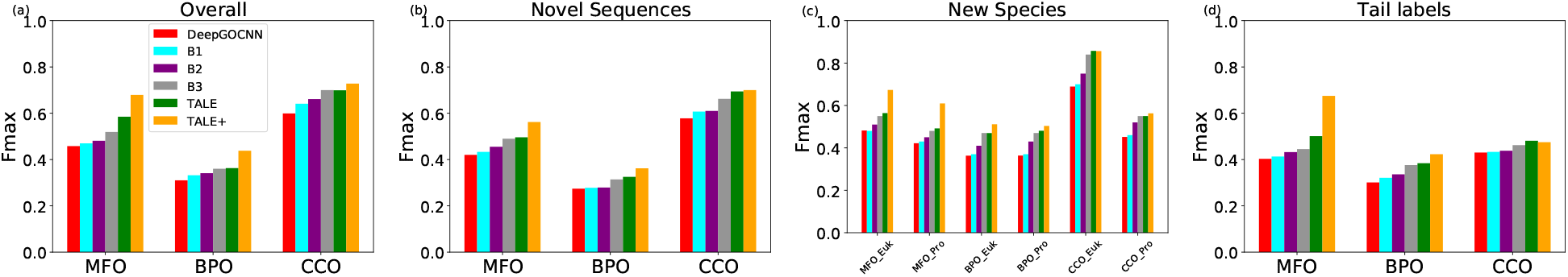
The Fmax performances of various models in the ablation study for (a) the overall test set as well as test sequences (b) with MSI≤30%, (c) in new eukaryotic or prokaryotic species and with MSI≤40%, or (c) label frequency ≤ 20%.

Various generalizability of the above models is also assessed over test sequences with MSI≤ 30%, in new species (and with MSI≤40%), or never or rarely (LF≤20%), as summarized in Fig. 5(b)–(d), respectively. Exact Fmax and AuPRC values and detailed analyses are in SI Sec. 8. Transformers and/or joint sequence–label embedding were the biggest contributors to nearly all AuPRC and most Fmax in all generalizability types and all ontologies considered. Sequence similarity often contributes significantly to generalizability (measured by Fmax) in MFO and occasionally in BPO.

## 4 Conclusion

In this paper, we have developed a novel transformer-based deep learning model named TALE, with joint embedding of sequence inputs and hierarchical function labels. The transformer architecture could learn sequence embedding while considering the long-term dependency within the sequence, which could generalize better to sequences with low similarity to the training set. To further generalize to tail labels (functions never or rarely annotated in the training set), we learn the label embedding, jointly with the sequence embedding, and use their joint similarity to measure the contribution of each amino acid to each label. The similarity matrix is further used to reweigh the contributions of each amino acid toward final predictions. In addition, we use TALE+, a convex combination of TALE and a similarity-based method, DIAMOND, to further improve model performances and generalizability.

Our results on a time-split test set demonstrate that TALE+ outperformed all sequence-based methods in all three ontologies and outperformed the state-of-the-art hybrid method (using network information) in BPO and CCO. When network information is not available, TALE+ outperformed all competing methods in all ontologies. Importantly, both TALE and TALE+ showed superior generalizability to sequences of low homology (and in never or rarely annotated species) and rarely annotated functions, echoing the rationales of our algorithm development. Ablation studies indicate that our newly introduced algorithmic components, especially transformer encoders and joint sequence–label embedding, contributed the most to such sequence, species, and function generalizability, whereas sequence similarity-based DIAMONDScore also helped.

Both TALE and TALE+ are fast models that can annotate 1,000 sequences within a couple of minutes on a mid-range GPU (Nvidia K80). These high-throughput annotators with both accuracy and generalizability would help close the increasing gap between high-throughput biological data and deep biological insights. In future, integrating additional data beyond protein sequences, particularly protein interaction networks, would further help close the gap especially for the ontologies of biological processes and cellular components.

## Acknowledgements

We thank Texas A&M High Performance Computing Resources for GPU allocations.

## Funding

This work has been supported by the National Institute of General Medical Sciences (R35GM124952).

## Notes

https://github.com/Shen-Lab/TALE

### Competing Interest Statement

The authors have declared no competing interest.

