## Supporting Information for "TALE: Transformer-based protein function Annotation with joint sequence–Label Embedding"

#### 1 Hyperparameters, training loss and validation accuracy

| ontology | MFO | | | BPO | | | CCO | | |
|---|---|---|---|---|---|---|---|---|---|
| models | model1 | model2 | model3 | model1 | model2 | model3 | model1 | model2 | model3 |
| $\lambda$ | 10 | 0.5 | 1.0 | 10 | 1.0 | 0.5 | 1.0 | 0.5 | 10 |
| learning rate | 0.001 | 0.001 | 0.001 | 0.001 | 0.001 | 0.001 | 0.001 | 0.001 | 0.001 |
| batch size | 32 | 32 | 32 | 16 | 16 | 16 | 32 | 32 | 32 |

Table S1: The hyperparameter settings of top 3 models in TALE. $\lambda$ was tuned on the grid {0.1, 0.5, 1.0, 2.0, 5.0, 10}. The learning rate was tuned on the grid {0.1, 0.01, 0.001, 0.0001}. The batch size was tuned on the grid {8, 16, 32, 64, 128}.

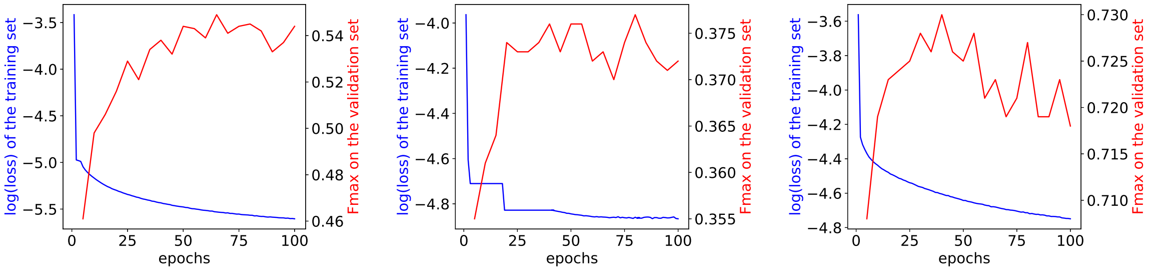


(a) MFO (b) BPO (c) CCO

Figure S1: The curves of the training loss and Fmax on the validation set of the best TALE model for (a): MFO, (b): BPO, (c): CCO.

### 2 The distributions of test sequences in Maximum Sequence Identity (MSI)

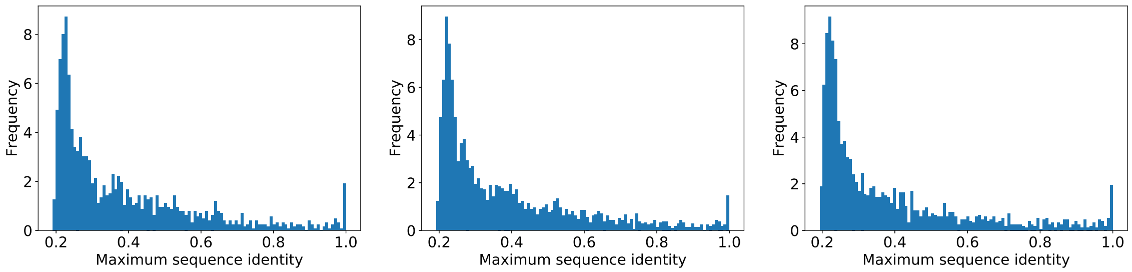


(a) (b) (c)

Figure S2: The distributions of maximum sequence identity between each test sequence and all training sequences for (a): MFO, (b): BPO, (c): CCO.

### 3 The distributions of test sequences in new species and low homology

#### 3.1 Statistics related to species

| Ontology | MFO | BPO | CCO |
|---|---|---|---|
| #Species in the training set | 1298 | 1346 | 757 |
| #Species in the test set | 333 | 314 | 164 |
| #Species in the test set but not the training set | 161 | 137 | 85 |
| #Samples in the test set with NSSS$\neq$0 | 1345 | 2374 | 1718 |
| #Samples in the test set with NSSS=0 | 214 | 236 | 189 |
| #Total samples in the test set | 1559 | 2610 | 1907 |

Table S2: The statistics of species over various data sets and the statistics of test sequences, with zero NSSS (new species) or not.

From Table S3, we found that around 48% (MFO), 44% (BPO), and 52% (CCO) of test species are new to the training set. We also found that around 14% (MFO), 9% (BPO), and 10% (CCO) of test sequences are with new species never seen in the training set.

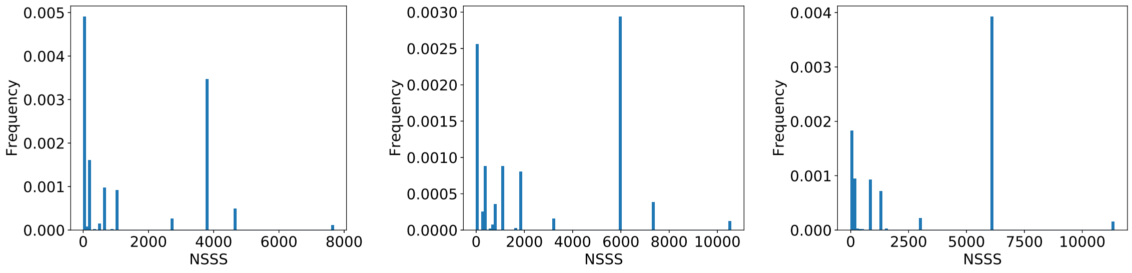


(a) (b) (c)

Figure S3: The distributions of NSSS for test sequences. (a): MFO, (b): BPO, (c): CCO.

Among the 214, 236, and 189 test sequences in new species for MFO, BPO, and CCO, respectively, we further filtered them based on maximum sequence identity (MSI) to the training set and only retained those with MSI within 40%. The statistics of resulting test sequences in new species and low homology are provided as follows.

| Domain/Ontology | MFO | BPO | CCO |
|---|---|---|---|
| Bacteria | 99 | 70 | 54 |
| Eukaryota | 53 | 80 | 86 |
| Archaea | 4 | 4 | 0 |
| Viruses | 4 | 6 | 5 |

Table S3: The statistics of test sequences in new species and low homology (MSI$\leqslant$ 40%) across 4 domains and 3 ontologies.

### 4 The distributions of test sequences in their label frequencies ($\mathrm{LF}(i)$) in the training set

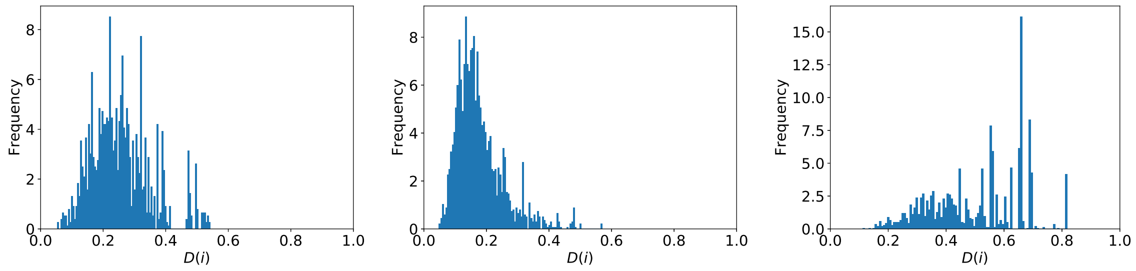


(a) (b) (c)

Figure S4: The histograms of test sequences' label frequencies. (a): MFO, (b): BPO, (c): CCO.

### 5 Model complexity

| | DeepGOPlus | TALE |
|---|---|---|
| # Trainable parameters | 54,585,424 | 21,813,855 |

Table S4: The numbers of trainable parameters for DeepGOPlus and TALE. For fair comparison, the number for TALE is the sum of 9 models (TALE uses top 3 models averaged for each of the three ontologies).

### 6 Generalizability test using equal-populated bins

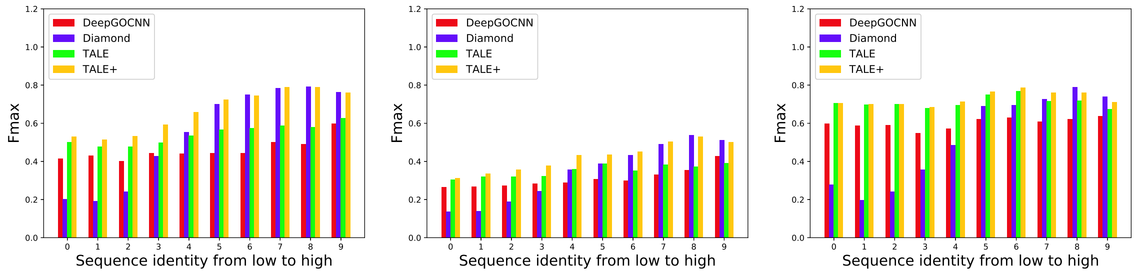


(a) MFO (b) BPO (c) CCO

Figure S5: The Fmax values for 10 test subsets of increasing sequence identity MSI.

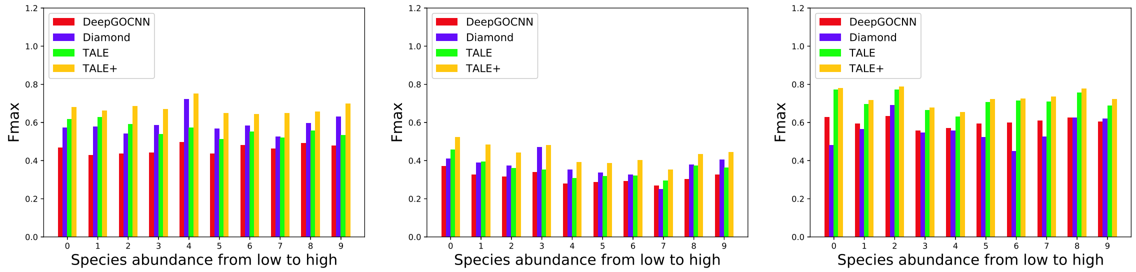


(a) MFO (b) BPO (c) CCO

Figure S6: The Fmax values for 10 test subsets of increasing species abundance. Note that bin 0 is the subset with NSSS=0, and bins 1-9 are equal-populated with the rest (NSSS≠0).

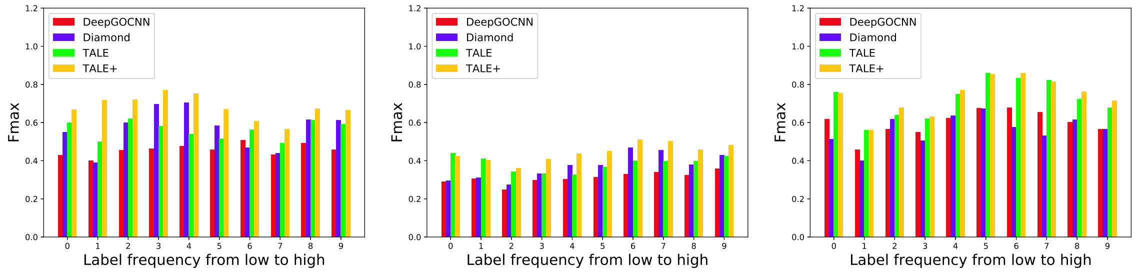


(a) MFO (b) BPO (c) CCO

Figure S7: The Fmax values for 10 test subsets of increasing label frequency LF(·).

### 7 Ablation study – Overall performance over the test set

We note that, from DeepGOCNN to TALE+, both Fmax and AuPRC are gradually increasing over model variants. Apparently each component of TALE and TALE+ gave non-trivial contributions to the overall performances, as summarized below for individual ontologies.

| | Fmax | | | AuPRC | | |
|---|---|---|---|---|---|---|
| Ontology | MFO | BPO | CCO | MFO | BPO | CCO |
| DeepGOCNN | 0.458 | 0.310 | 0.599 | 0.345 | 0.176 | 0.506 |
| **B1** (+transformer) | 0.470 | 0.332 | 0.641 | 0.433 | 0.226 | 0.626 |
| **B2** (+hierarchical reg.) | 0.481 | 0.341 | 0.661 | 0.444 | 0.248 | 0.652 |
| **B3** (+joint embedding) | 0.519 | 0.360 | 0.700 | 0.467 | 0.252 | 0.683 |
| TALE | 0.585 | 0.363 | 0.699 | 0.564 | 0.280 | 0.685 |
| TALE+ | **0.679** | **0.438** | **0.728** | **0.613** | **0.316** | **0.711** |

Table S5: The performances of DeepGOCNN (base), TALE+, and ablated TALE+ variants in between, over 1559, 2610, and 1907 test sequences in MFO, BPO and CCO, respectively.

In MFO, both transformers and label embedding (jointly with sequence embedding) steadily improved Fmax by 0.02 to 0.04. Using an ensemble of models in TALE saw a larger Fmax improvement over 0.06 whereas adding similarity-based DIAMOND had the largest Fmax improvement over 0.09. This echoes our hypothesis that sequence similarity carries much more information about biochemical functions compared to other ontologies. Of course, all components introduced to TALE together improved Fmax more by 0.13.
In BPO, based on Fmax, again transformers and label embedding steadily improved Fmax by 0.02 to 0.03. Ensemble average did not help Fmax noticeably whereas similarity-based DIAMONDScore again had the largerst Fmax boost of over 0.07.
In CCO, unlike MFO and BPO, the most important contributions actually came from using transformers and label embedding, each of which improved Fmax by 0.04 to 0.058. Whereas ensemble average did not improve Fmax here, similarity-based DIAMONDScore slightly improved Fmax by over 0.02.
We also note that the AuPRC performances had similar but not identical trends as the Fmax ones.

### 8 Ablation Study - Generalizability

In this last subsection of results, we perform ablation studies to assess the contributions of algorithmic components toward various generalizability (rather than overall performances discussed before). Therefore, we assess aforementioned model variants from DeepGOCNN to TALE+ over three subsets of the test set where generalizability is needed the most: sequences with low homology to the training set (maximum sequence identity [MSI] $\leqslant$ 30%), sequences of species never annotated (NSSS=0) and of low homology (MSI$\leqslant$40%) in the training set, and sequences of functions rarely annotated in the training set (average label frequency below 0.2). The ablation results toward sequence, species, and function generalizability are summarized in Tables S6, S7, and S8, respectively.

| | Fmax | | | AuPRC | | |
|---|---|---|---|---|---|---|
| Ontology | MFO | BPO | CCO | MFO | BPO | CCO |
| DeepGOCNN | 0.420 | 0.274 | 0.578 | 0.293 | 0.134 | 0.491 |
| **B1** | 0.433 | 0.278 | 0.607 | 0.314 | 0.167 | 0.514 |
| **B2** | 0.455 | 0.279 | 0.610 | 0.324 | 0.163 | 0.517 |
| **B3** | 0.490 | 0.314 | 0.662 | 0.378 | 0.207 | 0.645 |
| TALE | 0.496 | 0.325 | 0.694 | 0.429 | **0.224** | **0.686** |
| TALE+ | **0.562** | **0.362** | **0.700** | **0.439** | 0.217 | 0.680 |

Table S6: The performances of DeepGOCNN (base), TALE+, and ablated TALE+ variants in between, over 762, 1251, and 992 test sequences with maximum sequence identity below 0.3, in MFO, BPO and CCO, respectively.

Interestingly, compared to our earlier observations in Table S5, similarity-based DIAMONDScore was found not necessarily the largest single contributor even in MFO and BPO. Joint sequence–label embedding (and transformers) alone contributed the most in MFO when the test species were never annotated in the training

|  | Fmax |  |  |  |  |  |
|---|---|---|---|---|---|---|
|  | MFO |  | BPO |  | CCO |  |
| Ontology | eukaryotes | prokaryotes | eukaryotes | prokaryotes | eukaryotes | prokaryotes |
| DeepGOCNN | 0.482 | 0.422 | 0.363 | 0.364 | 0.689 | 0.451 |
| **B1** | 0.480 | 0.430 | 0.370 | 0.370 | 0.700 | 0.460 |
| **B2** | 0.510 | 0.450 | 0.410 | 0.430 | 0.750 | 0.520 |
| **B3** | 0.550 | 0.480 | 0.470 | 0.470 | 0.840 | 0.550 |
| TALE | 0.564 | 0.492 | 0.470 | 0.481 | **0.857** | 0.550 |
| TALE+ | **0.673** | **0.610** | **0.511** | **0.504** | 0.856 | **0.563** |
|  | AuPRC |  |  |  |  |  |
|  | MFO |  | BPO |  | CCO |  |
| Ontology | eukaryotes | prokaryotes | eukaryotes | prokaryotes | eukaryotes | prokaryotes |
| DeepGOCNN | 0.369 | 0.302 | 0.257 | 0.197 | 0.490 | 0.313 |
| **B1** | 0.390 | 0.350 | 0.262 | 0.230 | 0.534 | 0.356 |
| **B2** | 0.464 | 0.425 | 0.263 | 0.252 | 0.588 | 0.389 |
| **B3** | 0.502 | 0.452 | 0.289 | 0.358 | 0.673 | 0.442 |
| TALE | 0.578 | 0.476 | 0.302 | **0.381** | **0.775** | **0.485** |
| TALE+ | **0.618** | **0.520** | **0.335** | 0.355 | 0.746 | 0.460 |

Table S7: The performances of DeepGOCNN (base), TALE+, and ablated TALE+ variants in three ontologies, over eukaryotes and prokaryotes with NSSS=0 (new species) and MSI $\leqslant$ 40% (low homology)

.

|  | Fmax |  |  | AuPRC |  |  |
|---|---|---|---|---|---|---|
| Ontology | MFO | BPO | CCO | MFO | BPO | CCO |
| DeepGOCNN | 0.403 | 0.301 | 0.430 | 0.316 | 0.184 | 0.400 |
| **B1** | 0.413 | 0.321 | 0.433 | 0.456 | 0.204 | 0.454 |
| **B2** | 0.432 | 0.336 | 0.438 | 0.478 | 0.211 | 0.458 |
| **B3** | 0.445 | 0.376 | 0.462 | 0.545 | 0.260 | 0.478 |
| TALE | 0.501 | 0.384 | **0.481** | 0.580 | 0.273 | 0.496 |
| TALE+ | **0.675** | **0.423** | 0.475 | **0.630** | **0.314** | **0.515** |

Table S8: The performances of DeepGOCNN (base), TALE+, and ablated TALE+ variants in between, over 463, 1874, and 35 test sequences with LF($\cdot$) $\leqslant$ 0.2 (of labels rarely seen in the training set), in MFO, BPO and CCO, respectively.

set. In CCO, the largest single contributor to Fmax improvement was label embedding and/or transformers in all the three generalizability cases.

When we compare all components introduced to TALE together (from DeepGOCNN to TALE) with DIAMONDScore (from TALE to TALE+), their Fmax improvements were 0.076 vs. 0.066 in MFO, 0.051 vs. 0.037 in BPO, and 0.114 vs. 0.006 in CCO, when sequence identities are low. These improvements were 0.082 vs. 0.109 (0.070 vs. 0.118) in MFO, 0.107 vs. 0.041 (0.117 vs. 0.023) in BPO, and 0.168 vs. -0.001 (0.099 vs. 0.013) in CCO, when sequences are in never-annotated eukaryotic (prokaryotic) species and low homology compared to the training set. And they were 0.098 vs. 0.174 in MFO, 0.084 vs. 0.039 in BPO, and 0.051 vs. -0.006 in CCO, when functions are rarely annotated in the training set. The trends are even more clear for AuPRC. Therefore, in all cases, we found that TALE components collectively improved model generalizability more, although similarity-based DIAMONDScore also contributed to Fmax (except in CCO).
